# Disruption of Marrow Microenvironments in Chronic Lymphocytic Leukemia by High-Resolution Synchrotron Micro-Computed Tomography

**DOI:** 10.1101/2025.10.20.683519

**Authors:** In Kyu Lee, Yoshihiro Obata, Anthony D. Pomicter, Justin A. Williams, Bejal Kikani, Christopher E. Jensen, Douglas W. Sborov, Deborah M. Stephens, Claire Acevedo

## Abstract

Chronic lymphocytic leukemia (CLL) is associated with increased fracture risk unexplained by standard bone density scans, suggesting underlying microstructural alterations. To investigate this, we used high-resolution synchrotron micro-computed tomography (SRµCT) on bone marrow biopsies from 17 CLL patients, who were stratified into low and high infiltration groups using objective, data-driven clustering. To our knowledge, this is the first report to quantify these changes.

High CLL marrow infiltration was associated with a 40.3% reduction in lacuna density and a 101% increase in the normalized adipose surface area-to-volume ratio, a metric indicating greater structural fragmentation. Both changes correlated with leukemic infiltration percentage and showed partial reversal after therapy in a longitudinal case. Furthermore, high CLL burden significantly altered the morphological distributions of the remaining lacunar osteocyte (p *<* 0.001). We identify a novel marrow remodeling phenotype in CLL characterized by osteocyte depletion and adipose disruption. These changes likely contribute to skeletal fragility and represent potential microstructural biomarkers for assessing marrow health more accurately than conventional imaging.

## Introduction

Chronic lymphocytic leukemia (CLL) is the most prevalent adult leukemia in Western countries, with over 200 000 currently living with CLL in the United States alone [1]. CLL is characterized by an accumulation of neoplastic B-lymphocytes in the peripheral blood, bone marrow, and lymphoid tissues. CLL typically presents at an older age with a median age at diagnosis of 70 years [2]. While there is a demographic overlap in the populations at risk for both CLL and osteoporosis [3], CLL has been independently associated with lower bone density and increased incidence of fragility fractures. Importantly, CLL has been linked to axial skeleton fractures, which can lead to significant morbidity and mortality [4]. Review of the SEER-Medicare database revealed that patients with CLL had a significantly increased risk of both vertebral (HR, 1.29, CI, 1.16–1.44, *P <*0.001) and pelvic fractures (HR, 1.56, CI, 1.33–1.83, *P <*0.001) compared to age-matched controls. In a previous study [5], we reviewed a group of patients with CLL and a diagnosis of osteoporosis or osteopenia and found a very high rate of fragility fractures (66%), and of those, 81% of the fractures were in the vertebrae. These fractures were present despite most patients having a non-osteoporotic T-score (*>*-2.5) on Dual-energy X-ray (DXA) scans, indicating that DXA scans are not a sufficient imaging modality to accurately predict a patient’s risk of axial fragility fractures [6].

Despite the breadth of existing and ongoing research in the field of CLL [7–12], there has been minimal attention directed toward how malignant cells physically reshape the bone marrow microenvironment, especially with regard to structural changes in bone and adipose tissues. In contrast, extensive research in other hematologic malignancies, such as multiple myeloma (MM), acute myeloid leukemia (AML), and acute lymphoblastic leukemia (ALL), provide clear precedents of tumor-induced alterations in osteocytes, osteoblasts, and bone marrow adipocytes that fuel disease progression and skeletal complications [13–15]. Whether comparable structural and compositional changes occur within the bone marrow microenvironment in CLL remains poorly established [16], underscoring a significant gap in clinical knowledge regarding fracture risk prediction and skeletal health management. Micro-computed tomography (microCT) has been instrumental in these contexts, capturing extensive bone loss, altered trabecular architecture, and marrow infiltration associated with malignant cell proliferation [17, 18].

High resolution synchrotron radiation microCT imaging (SRµCT) is an innovative technology that can be used to investigate the three-dimensional microstructure of the bone marrow and tissue composition. High-resolution imaging is critical for resolving individual osteocyte lacunae — small cavities that house osteocyte cells, which are responsible for mechanosensing as well as orchestrating local bone remodeling [19, 20]. In addition, imaging of both soft (adipose) and hard (bone) tissues simultaneously requires a technique capable of differentiating these materials despite their different x-ray absorption characteristics. Thus, a central technical challenge lies in segmenting bone tissue and adipose tissue within a single dataset, given the wide dynamic range of absorption signals. To address this challenge, our approach combines deep learning segmentation methods and builds upon recent developments in multi-tissue SRµCT imaging [21]. By enabling precise quantitative measurements of the bone and adipose microarchitecture, this pipeline offers a view of potential crosstalk among malignant B-cells, osteocytes, and marrow adipocytes (fat cells), similar to interactions observed in other blood cancers.

The novelty of the current study lies in three primary aspects. First, we collected bone marrow core biopsy samples from patients with CLL, allowing for direct observation of structural and compositional changes associated with leukemic infiltration. Second, our study employs a methodological approach combining high-resolution SRµCT imaging and deep learning segmentation algorithms. This integration enables us to simultaneously resolve detailed structural features of both bone and adipose tissues within a single dataset — overcoming previously encountered technical limitations. Third, we identified a previously unrecognized adipose-associated pathology specific to CLL, characterized by significant structural disruption in bone marrow adipose tissue correlated with leukemic burden.

In this study, we employed SRµCT to examine bone marrow biopsy samples from patients with varying levels of CLL involvement. Our primary aim was to examine whether the extent of leukemic infiltration correlates with osteocyte lacuna and adipose microarchitecture. Through these novel insights, we aim to establish new microstructural biomarkers and identify targets for interventions aimed at mitigating highly morbid fragility fractures in CLL patients.

## Materials and methods

### Patients and Sample Collection

Eighteen bone marrow core biopsy samples were collected from seventeen patients diagnosed with CLL (Table 1) who underwent bone marrow biopsy through iliac crest puncture as part of routine clinical care. An illustration of the biopsy procedure is shown in Fig. 1 A. One patient provided both pre-treatment and post-treatment samples. These samples were collected between November 2021 and September 2022 after obtaining informed patient consent, under an approved Institutional Review Board protocol (IRB# 00045880).

**Table 1.** Clinical Characteristics of Patients with CLL.

| Characteristic | Hi-CLL<br>(N=7) | Lo-CLL<br>(N=11) | Overall<br>(N=18) |
| --- | --- | --- | --- |
| Median % Marrow Core Involvement,<br>% (range) | 75 (40–99) | 0 (0–3) | 0.6 (0–99) |
| Median Patient Age at Sample<br>Collection, years (range) | 70 (68–85) | 69 (63–73) | 70 (63–85) |
| Median Time from CLL Diagnosis<br>to Sample Collection, years (range) | 3 (3–12) | 2 (1–15) | 3 (1–15) |
| Samples from Male Patient, number (%) | 3 (43) | 8 (73) | 11 (61) |
| Median Body Mass Index (BMI) at<br>Sample Collection, kg/m<sup>2</sup> (range) | 29 (22–37) | 29 (19–41) | 29 (19–41) |
| IGHV unmutated, number (%) | 4 (57) | 10 (91) | 14 (78) |
| **FISH\*** |  |  |  |
| Deletion 13q | 4 (57) | 7 (58) | 11 (61) |
| Trisomy 12 | 3 (43) | 5 (45) | 8 (44) |
| Deletion 11q | 3 (43) | 3 (25) | 6 (33) |
| Deletion 17p | 3 (43) | 2 (18) | 5 (28) |
| TP53 mutation, number (%) | 2 (29) | 1 (13) | 3 (17) |
| Complex karyotype ($\geq 3$ chromosomal<br>aberrations), number (%) | 4 (57) | 2 (18) | 6 (33) |
| Number of Lines of CLL-Directed Treatment<br>Prior to Sample Collection, number (range) | 0 (0–3) | 1 (1–6) | 1 (0–6) |
| **Patient Received the Following CLL-Directed Treatments\*** |  |  |  |
| Chemoimmunotherapy | 2 (29) | 1 (13) | 3 (17) |
| Bruton's Tyrosine Kinase Inhibitor | 1 (14) | 9 (82) | 10 (56) |
| Venetoclax | 2 (29) | 6 (55) | 8 (44) |
| **Types of CLL-Directed Treatments Patient Received Most Recently\*** |  |  |  |
| Chemoimmunotherapy | 1 (14) | 0 (0) | 1 (5) |
| Bruton's Tyrosine Kinase Inhibitor | 1 (14) | 10 (83) | 11 (58) |
| BCL2 Inhibitor | 1 (14) | 5 (42) | 6 (32) |
| Median Time from Most Recent CLL-Directed<br>Treatment to Sample Collection, months (range) | 0 (0–26) | 0 (0–4) | 0 (0–26) |

\*Not mutually exclusive

**Fig. 1.**
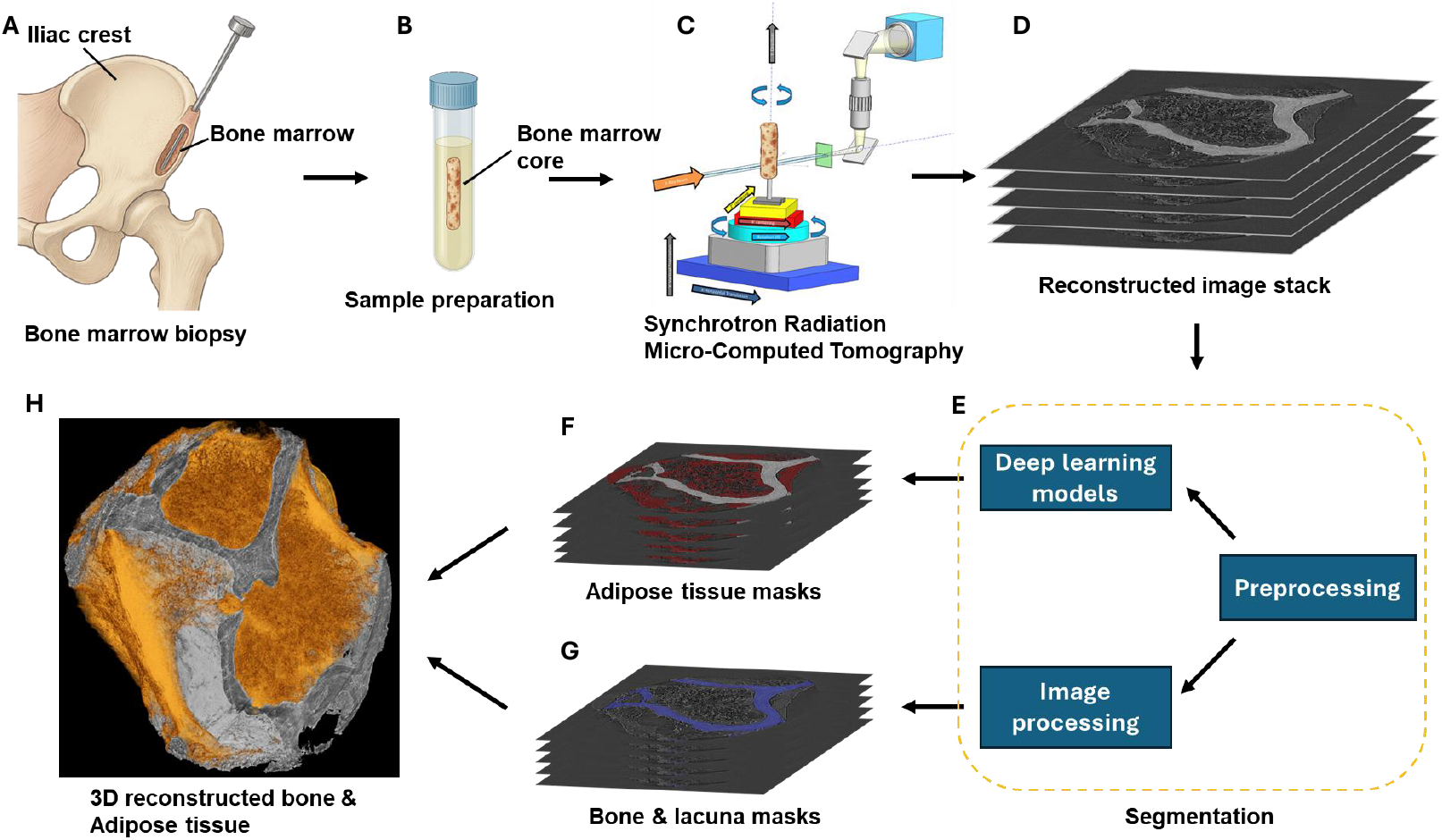
Overview of the imaging and segmentation pipeline. (A) Illustration of the bone marrow biopsy procedure performed at the iliac crest. (B) Extracted core biopsy sample stored in 70% ethanol at 4°C. (C) Imaging of the sample using synchrotron radiation micro-computed tomography (SRµCT).(D) Example of a reconstructed 2D image slice from the SRµCT dataset. (E) Image analysis pipeline, including preprocessing (8-bit conversion, intensity normalization), deep learning–based segmentation of adipose tissue, and image processing–based segmentation of bone and osteocyte lacunae using Otsu thresholding, morphological operations, and size filtering. (F) Output of adipose tissue segmentation. (G) Output of bone and lacuna segmentation. (H) 3D reconstruction showing segmented bone and adipose tissue architecture.

Bone marrow core biopsies were obtained using standard clinical biopsy needles and measured approximately 2 mm in diameter and 5 mm in length. Immediately following collection, cores were placed in 2 mL of 70% ethyl alcohol, 30% water and stored at 4°C (Fig. 1 B). The extent of CLL involvement in the bone marrow core biopsy specimens was determined through local pathology review. These assessments quantify the proportion of leukemic B-cells within the total marrow cellularity.

### Grouping strategy and unsupervised clustering

To objectively stratify patients into groups based on disease burden, we performed an unsupervised, one-dimensional k-means clustering of the percent marrow CLL involvement with k=2 specified a priori for clinical interpretability. Clustering was implemented in scikit-learn (k-means++ initialization; multiple random starts; fixed random seed for reproducibility). This resulted in two distinct cohorts for comparative analysis (Fig. 2 A): a Low CLL Involvement group (Lo-CLL, ≤3%, n=11) and a High CLL Involvement group (Hi-CLL, ≥40%, n=7). The quality of this clustering was quantified by the silhouette score, and the implied decision boundary was calculated as *t* = (*µ*_1_ + *µ*_2_)*/*2, where *µ*_1_and *µ*_2_ are the two cluster centroids.

**Fig. 2.**
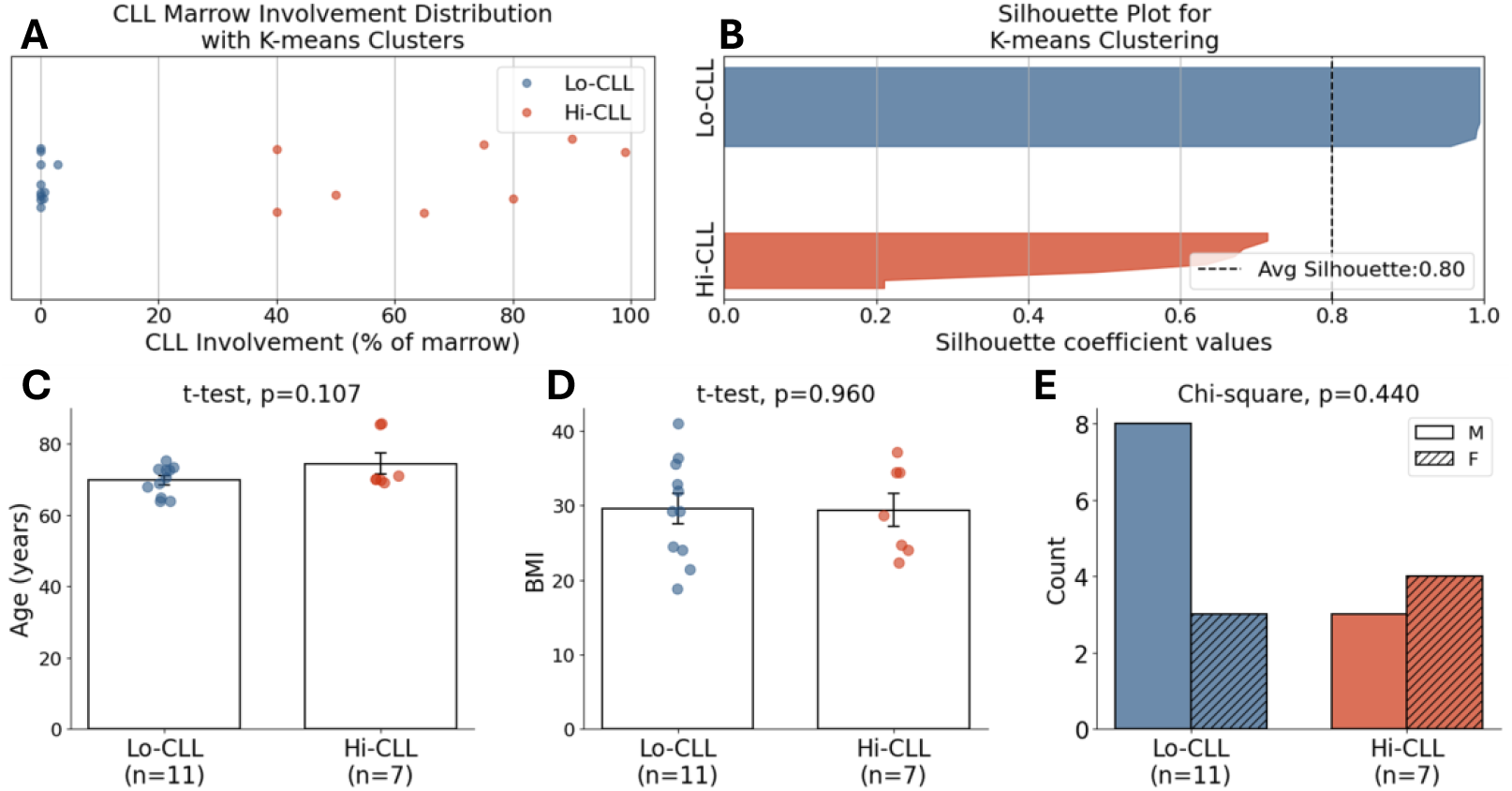
Data-driven patient stratification and clurstering group characteristics (A) Unsupervised k-means clustering of marrow CLL involvement percentage for all 18 patient samples. The analysis identified two distinct, non-overlapping clusters: Low CLL involvement (Lo-CLL, n=11, blue) and High CLL involvement (Hi-CLL, n=7, red). (B) Silhouette plot validating the robustness of the k-means clustering. All samples show a positive silhouette score, and the high average score (S=0.80, black dashed line) confirms a strong separation between the two clusters. (C-E) Comparison of base-line demographic and clinical characteristics between the Lo-CLL and Hi-CLL groups. No statistically significant differences were observed in (C) patient age, (D) Body Mass Index (BMI), or (E) sex distribution.

### Synchrotron radiation micro-computed tomography (SRµCT)

SRµCT imaging was performed at beamline 8.3.2 at the Advanced Light Source in Berkeley, CA. Bone marrow cores contained both bone marrow adipose tissue and trabecular bone. Images were acquired with a beam energy of 22 keV and a 100 ms exposure time. The imaging setup is illustrated in Fig. 1 C. During imaging, a total of 1969 projection angles were acquired over a 180° sample rotation. Following image acquisition, reconstruction of the SRµCT datasets was conducted using the open-source Python library TomoPy [22], employing a filtered back-projection algorithm optimized for synchrotron data. The final reconstructed voxel size was 1.6 µm, ensuring sufficient spatial resolution to resolve both bone and adipose microstructures.

### Image Segmentation

High-resolution SRµCT datasets enabled segmentation and quantification of three key components of the marrow microenvironment: mineralized bone, osteocyte lacunae, and adipose tissue. Due to distinct differences in X-ray attenuation and morphology, each tissue compartment was segmented using a tailored approach combining classical image processing and deep learning methods.

#### Bone Segmentation

Bone tissue was segmented by leveraging its high X-ray attenuation relative to surrounding structures. First, the reconstructed SRµCT images were converted to 8-bit grayscale to standardize voxel intensity across datasets. We then applied Otsu’s thresholding [23], a histogram-based method that selects an optimal global threshold value by minimizing intra-class variance. This threshold separates the image into two classes—low-intensity voxels (background, soft tissue) and high-intensity voxels (mineralized bone).

Only voxels above the Otsu-derived threshold were retained, forming an initial binary mask of high-density mineralized bone. Because noise and artifacts can introduce small, spurious bright regions, a 3D morphological opening was applied to remove disconnected high-intensity specks. To isolate the contiguous bone sample from other segmented structures (e.g., debris or imaging artifacts), we computed the volume of all connected components and preserved only the largest component, which corresponded to the true bone core.

However, this initial mask often excluded internal voids such as osteocyte lacunae and vascular canals due to their lower intensity. To restore these biologically important internal features, we applied a 3D morphological closing, which fills small holes within the segmented structure, ensuring a complete and contiguous bone mask suitable for lacuna analysis.

#### Osteocyte Lacuna Segmentation

After the bone region was fully segmented, we proceeded to isolate osteocyte lacunae, which appear as small, ellipsoidal low-density voids embedded within the mineralized matrix. Because these features are defined by their contrast relative to surrounding bone, we performed binary thresholding within the previously defined bone mask to identify dark regions indicative of lacunar cavities. To ensure that only internal voids were included—excluding surface artifacts or external cavities—we first eroded the bone mask using a 3D structuring element. This erosion operation created a conservative mask that excluded boundary regions where partial volume effects and edge artifacts are more prevalent. We then inverted this eroded mask and removed any segmented cavities that touched it, effectively filtering out lacuna-like structures not fully embedded in bone.

Following this, all remaining cavity candidates were evaluated based on their volume. We retained only those within the biologically plausible range of 50 to 1000 µm^3^, as previously reported in human bone studies [24, 25]. This step removed small features likely to be noise and larger structures (such as vascular canals) that do not represent true lacunae. The final lacuna segmentation outputs were then used for downstream morphometric analyses, including lacunar density, volume, and shape, providing a quantitative assessment of osteocyte lacunae organization in relation to CLL infiltration.

#### Adipose Tissue Segmentation

Unlike mineralized bone, which exhibits high X-ray attenuation, bone marrow adipose tissue appears as a low-density, radiolucent structure in SRµCT images. However, the voxel intensity of adipose tissue often overlaps substantially with background noise and non-tissue regions, making traditional thresholding techniques unreliable for accurate segmentation. To address this challenge, we implemented a deep learning–based approach using convolutional neural networks (CNNs) to identify and delineate adipose tissue regions with higher specificity.

Representative 2D slices from the midsection of each 3D SRµCT volume were manually annotated to serve as ground truth labels. These annotated slices were used to train two CNN architectures for segmentation: U-Net and U-Net++ [26, 27]. To enhance the generalizability of the models, we applied data augmentation, including random horizontal and vertical flips, rotation between –90° and 90°, and zoom scaling from 0.5 to 1.5.

Both models were trained using a combined loss function, consisting of a weighted sum of Dice loss and cross-entropy loss (0.5 each), which balances pixel-wise classification accuracy with segmentation overlap performance. Optimization was performed using the Adam optimizer [28] with a weight decay of 1×10^3^ for 100 epochs.

At inference time, we implemented an ensemble strategy by averaging the probability maps produced by both models, improving segmentation robustness against inter-sample variability. Final adipose tissue masks were then manually reviewed and corrected where necessary to ensure anatomical accuracy and consistency across scans. This deep learning pipeline enabled reliable quantification of adipose morphology within the marrow space and facilitated comparative analysis between Lo-CLL and Hi-CLL patient groups. Notably, it allowed for the identification of adipose fragmentation patterns, such as irregular borders and disrupted continuity, that were not detectable via simple intensity-based methods.

### SRµCT data analysis

Several quantitative parameters were extracted from segmented SRµCT images to characterize bone microstructure. Within the segmented bone regions, we computed the tissue mineral peak density, defined as the most frequent intensity value (mode) within the tissue mineral density histogram. This metric reflects the predominant density level of the mineralized bone matrix, providing insight into bone quality and mineralization patterns.

Osteocyte lacuna characteristics were also quantified following segmentation. A 3D connected component analysis was utilized to identify and quantify individual osteocyte lacunae. Lacuna density was subsequently calculated as the number of osteocyte lacunae per unit bone volume. To assess the properties of the lacunar population, the volume and aspect ratio (ratio between the major and minor axes) were calculated for every individual lacuna. For each patient sample, the median lacunar volume and aspect ratio were determined, and these patient-level medians were then compared between the Lo-CLL and Hi-CLL groups. In addition, to compare the overall distributions more comprehensively, all individual lacunae from patients within each cohort were pooled, and a two-sample Kolmogorov-Smirnov test was used to detect significant differences in the distributions of lacunar volume and aspect ratio.

Adipose tissue parameters were obtained after robust segmentation using deeplearning methods. Adipose density (g/cm^3^) was calculated from raw SRµCT intensity values calibrated using the Center for X-ray Optics Database [29] to ensure accurate density measurements. Additionally, adipose tissue fragmentation was assessed using a normalized adipose tissue surface area-to-volume ratio (SA:V). As the standard SA:V ratio can be artificially inflated in small volumes due to scale effects, we employed a normalized ratio (*SA/V* ^2*/*3^) to produce a scale-invariant, unitless metric. This normalized SA:V effectively captures the degree of adipose tissue fragmentation, enabling reliable comparisons across samples of varying adipose tissue volumes.

### Statistical Analysis

Based on the cluster assignments, group comparisons of SRµCT variables were performed using Student’s t-tests. Pearson’s correlation analysis was employed to evaluate relationships between the percentage of CLL marrow involvement (as a continuous variable) and each SRµCT feature. Chi-squared tests were used to assess differences in categorical variables, such as sex distribution between groups. All statistical analyses were conducted using a SciPy library version 1.14.1 [30] in Python 3.12.7. Statistical significance was defined as a two-tailed *P <* 0.05.

## Results

### Data-driven partition of marrow involvement

To objectively stratify patients, unsupervised k-means clustering was performed on the percentage of marrow CLL involvement. The analysis partitioned the 18 patient samples into two distinct, non-overlapping cohorts: a Low CLL involvement group (Lo-CLL; 0–3%, cluster centroid at 0.38%) and a High CLL involvement group (Hi-CLL; 40–99%, cluster centroid at 67.4%) (Fig. 2A). The robustness of this datadriven partition was confirmed by strong cluster separation metrics (silhouette score S=0.80, Calinski–Harabasz index CH=99.72), visually represented by the silhouette plot (Fig. 2B). Furthermore, the resulting Lo-CLL and Hi-CLL cohorts were well-matched, showing no statistically significant differences in age, BMI, or sex (Fig. 2, panels C-E). This objective stratification provides a validated basis for the comparative analyses presented in the subsequent results.

### Osteocyte Lacunar Density was Decreased by 40% with High CLL Involvement

Qualitative assessment demonstrated structural differences in osteocyte lacunae between the two groups (Fig. 3, panels A–D). Lacunae appeared more densely and uniformly distributed within Lo-CLL samples (Fig. 3, panels A–B), whereas Hi-CLL samples exhibited fewer, sparsely distributed lacunae (Fig. 3, panels C–D). Quantitatively, lacuna density was significantly lower in the Hi-CLL group (17 100 lacunae per unit bone volume) compared to the Lo-CLL group (25 664 lacunae per unit bone volume), reflecting a 40.3% decrease (p *<* 0.05, Fig. 3E). Additionally, a significant negative correlation was identified between lacuna density and percentage of CLL infiltration (r = –0.49, p *<* 0.05, Fig. 3 F), strongly linking increased leukemic burden with decreased osteocyte lacunar presence. Notably, variance was greater in the Lo-CLL group, likely reflecting heterogeneity in disease duration and treatment exposure, yet the between-group difference and correlation in lacuna density remained statistically significant.

**Fig. 3.**
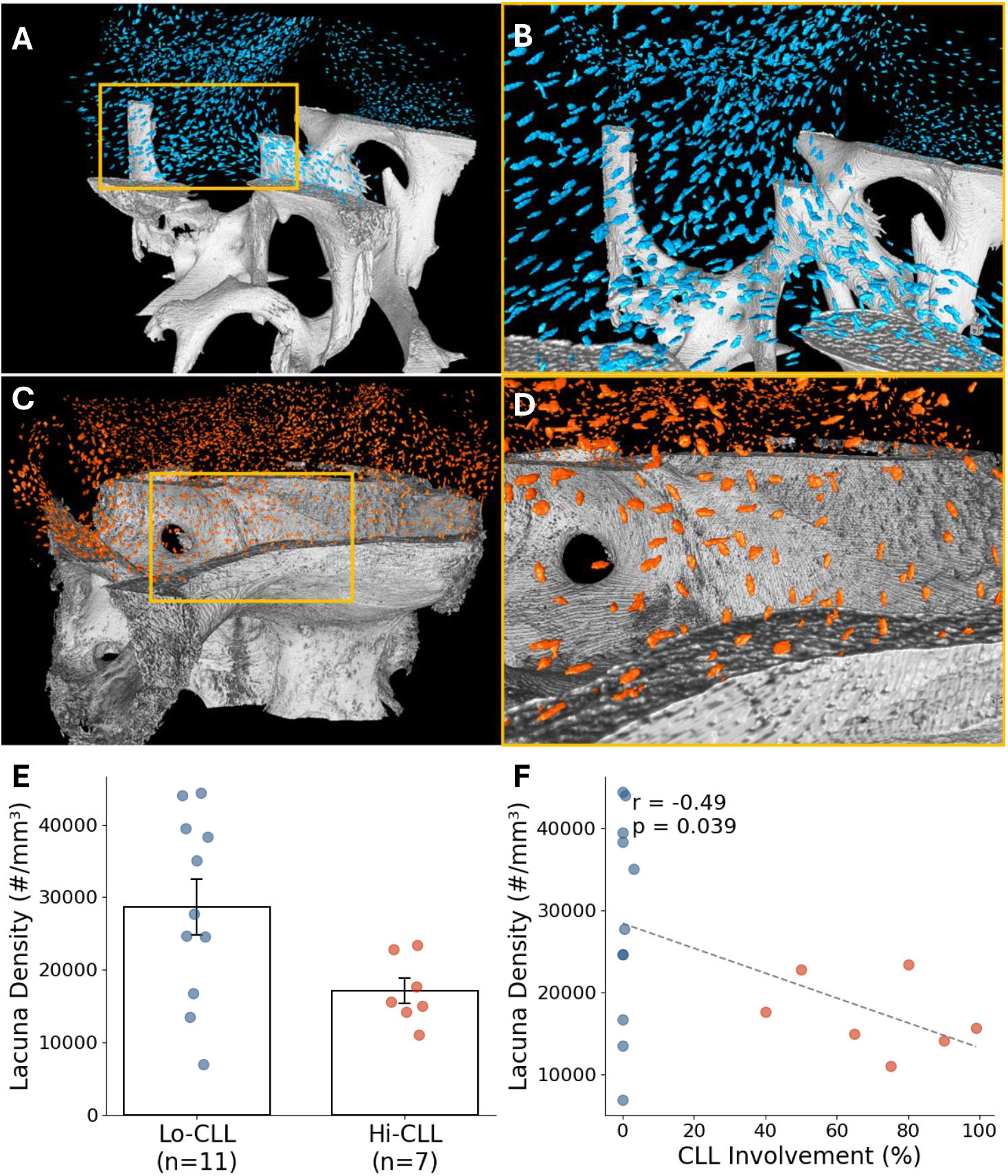
Lacunar density comparison between low and high CLL involvement groups. (A–D) Visualization of osteocyte lacunae in representative samples from low CLL involvement (A, B) and high CLL involvement (C, D) groups. Panels B and D are magnified views of the boxed regions in panels A and C, respectively. For scale, bone marrow cores were approximately 2 mm in diameter. (E) Quantitative comparison demonstrating 40.3% reduced lacuna density in high versus low CLL groups (p *<* 0.05). (F) Correlation plot illustrating a significant negative correlation between CLL involvement percentage and lacuna density (r = -0.49, p *<* 0.05).

### Bone Marrow Adipose Tissue Fragmentation was Increased by Two-fold with High CLL Involvement

Adipose tissue segmentation indicated significant fragmentation in samples from the Hi-CLL group compared to the Lo-CLL group. Representative images (Fig. 4, panels A–D) illustrate that adipose tissue within Lo-CLL samples (Fig. 4, panels A–B) appeared intact and smooth, while Hi-CLL samples (Fig. 4, panels C–D) displayed fragmented and irregular adipose structures. Additional examples from both Lo-CLL and Hi-CLL groups are provided in Supplementary Fig. 1, further demonstrating this pattern across multiple cases.

**Fig. 4.**
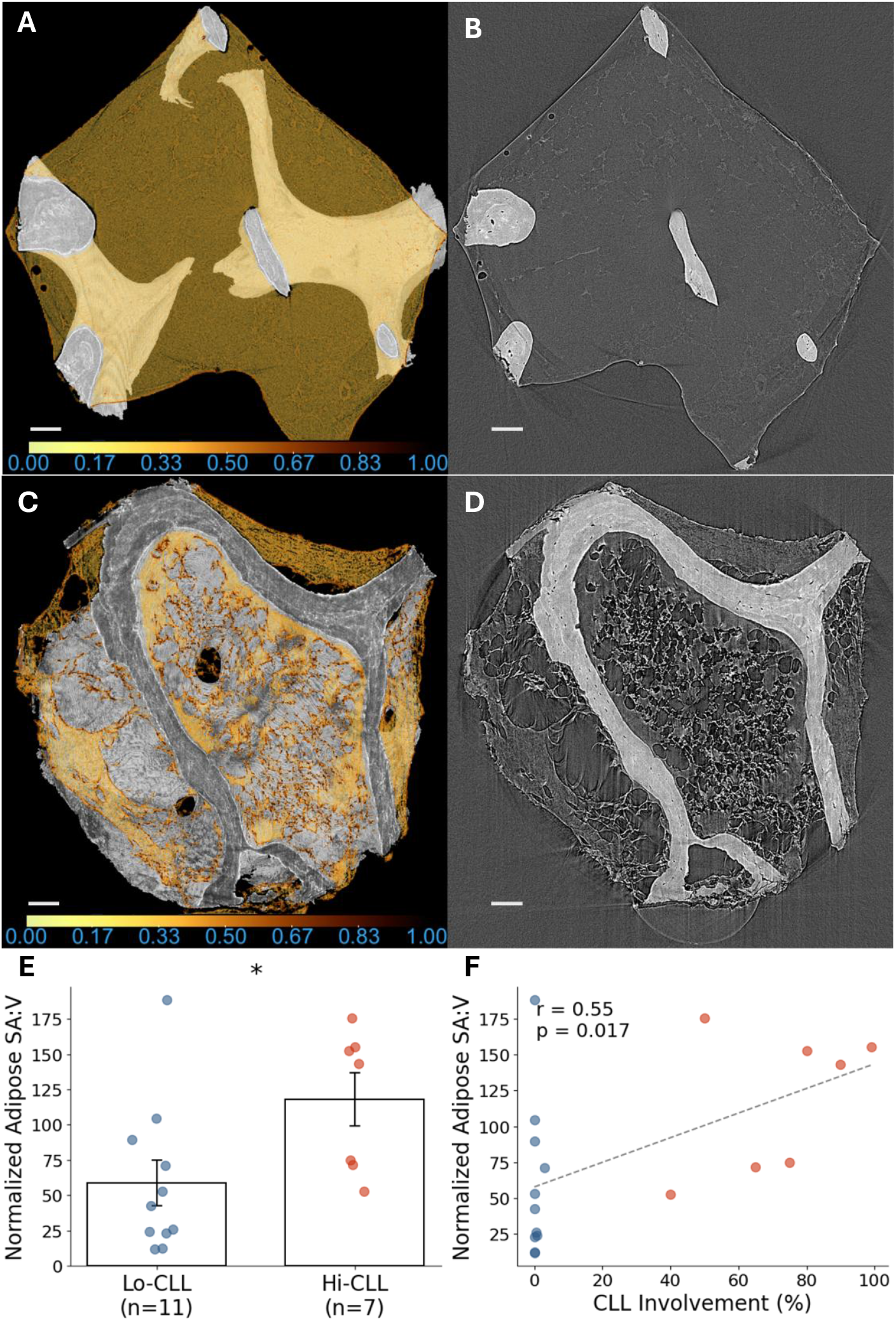
Structural disruption of adipose tissue with increasing CLL infiltration. (A, C) 3D visualizations of adipose tissue (yellow) within the bone marrow, with adipose density indicated by a normalized colormap. Brighter intensities correspond to higher adipose tissue density, while darker colors indicate lower density. (A, B) represent a Lo-CLL sample, while (C, D) represent a Hi-CLL sample. (B, D) Corresponding cross-sectional SRµCT images illustrate intact and smooth adipose architecture in the Lo-CLL sample (A, B), contrasting with fragmented and irregular adipose structures in the Hi-CLL sample (C, D). (E) Quantitative analysis reveals a 101% increased normalized adipose tissue SA:V in the Hi-CLL compared to the Lo-CLL. (F) The correlation plot shows a significant positive association between CLL involvement and normalized adipose SA:V. Scale bars, 100µm.

Quantitative analysis demonstrated significantly higher normalized adipose SA:V in the Hi-CLL group (118.0) compared to the Lo-CLL group (58.7), indicating a 101% increase in adipose fragmentation (p *<* 0.05, Fig. 4E). Moreover, normalized adipose SA:V correlated positively and significantly with the percentage of CLL involvement (r = 0.56, p *<* 0.05, Fig. 4 F). Importantly, mean BMI values were similar between the two groups, suggesting that adipose fragmentation observed at the microstructural level is more likely associated with CLL infiltration rather than overall body composition.

### Therapy-Induced Restoration of Marrow Microarchitecture in a Longitudinal Case Study

A single patient (male, 70 y, BMI = 24 kg/m^2^) contributed matched iliac-crest core biopsies obtained before and after ibrutinib therapy. The baseline specimen belonged to the Hi-CLL group (CLL involvement=65%), whereas the post-treatment specimen, with CLL involvement reduced to *<*1%, re-classified into the Lo-CLL group. Representative SRµCT cross-sections are shown in Fig. 5A–B: panel A (Hi-CLL) reveals numerous fragmented, serrated adipose profiles interspersed throughout the trabecular compartment, while panel B (Lo-CLL, post-therapy) shows a marked restoration of contiguous, smoothly contoured adipose tissue. We discovered a 194% increase in lacunar density (from 14 965 to 43 999 lacunae/mm^3^) and a 67% decrease in adipose tissue fragmentation, as measured by normalized surface area-to-volume ratio (from 71.7 to 24.0) after treatment. These paired changes are illustrated in Fig. 5C–D. No formal statistical testing was performed for this *n*=1 comparison; nevertheless, the paired observations illustrate that pharmacological reduction of leukemic burden is accompanied by large-magnitude improvements in both osteocyte and adipose microarchitecture.

**Fig. 5.**
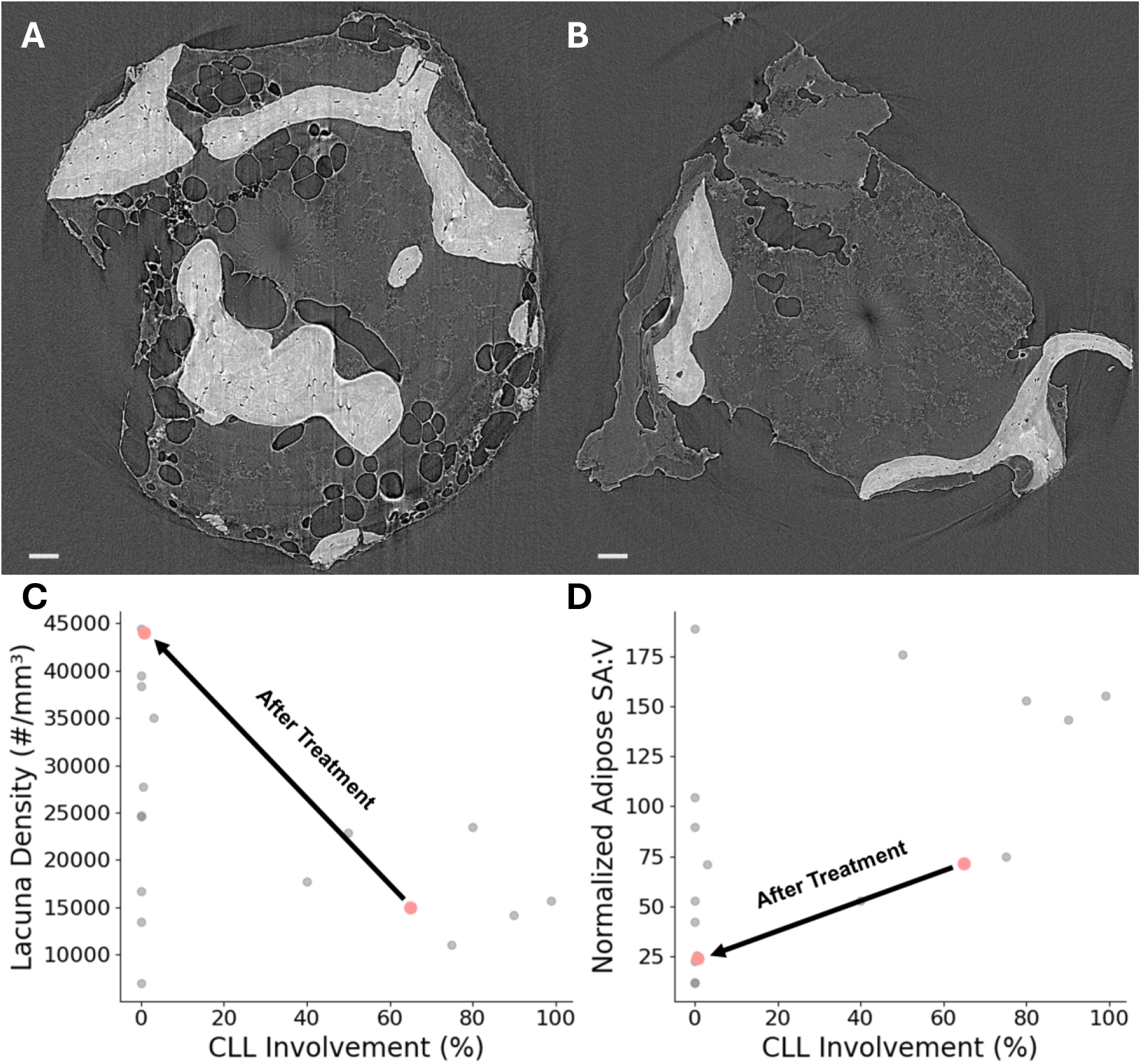
Pre- and post-treatment SR*µ*CT images and quantitative recovery of bone marrow microstructural parameters in a patient with CLL. (A) Cross-sectional 2D SR*µ*CT image from the patient before treatment, illustrating significant adipose tissue fragmentation. (B) Corresponding SR*µ*CT image from the same patient after treatment, showing a notable reduction in adipose fragmentation. Quantitative comparisons for osteocyte lacuna density (C) and normalized adipose SA:V (D)are shown with gray points representing other patients, while red points indicate this patient at two different time points (before and after treatment). Following treatment and the subsequent marked reduction in CLL marrow involvement, osteocyte lacuna density significantly increased, and normalized adipose SA:V substantially decreased, highlighting structural recovery within the bone marrow microenvironment.

### Increased Lacunar Volume is Correlated with Adipose Tissue Changes and Reduced Lacunar Density

Additional SRµCT-derived parameters provided further insights into the marrow microenvironmental changes associated with CLL. Correlation analyses (Fig. 6A) revealed significant relationships among these microstructural features. Median lacunar volume exhibited moderate positive correlations with tissue mineral peak intensity (r = 0.50) and adipose density (r = 0.70), suggesting interconnected alterations in bone mineralization and adipose tissue properties. Conversely, median lacunar volume negatively correlated with lacunar density (r = –0.51), reinforcing the idea that as osteocyte density declines, the remaining lacunae may become larger, possibly reflecting a compensatory structural adaptation or impaired osteocyte viability.

**Fig. 6.**
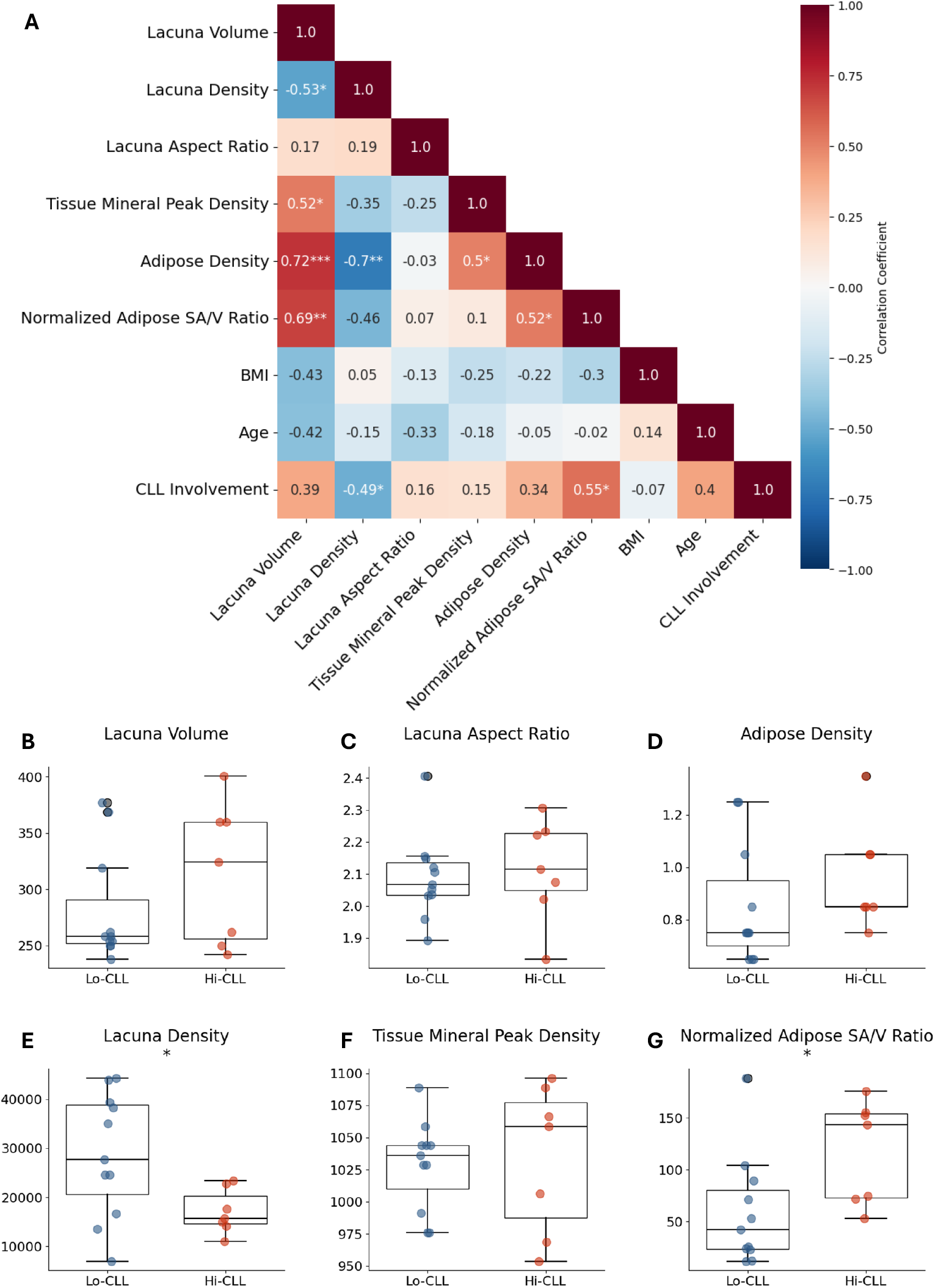
Bone and adipose microstructural alterations in CLL-infiltrated marrow. (A) Correlation matrix showing relationships between microstructural parameters and CLL involvement. Color intensity represents correlation coefficient (blue: negative, red: positive). Numbers indicate Pearson correlation coefficients. (B-G) Box plots comparing microstructural parameters between Lo-CLL and Hi-CLL. Parameters include lacunar volume, lacunar density, lacunar aspect ratio, tissue mineral peak density, adipose density, and normalized adipose surface area-to-volume ratio. Boxes show median and interquartile range, whiskers extend to 1.5× interquartile range, and individual points represent each sample. \**P <* 0.05, \*\**P <* 0.01, \*\*\**P <* 0.001.

**Fig. 7.**
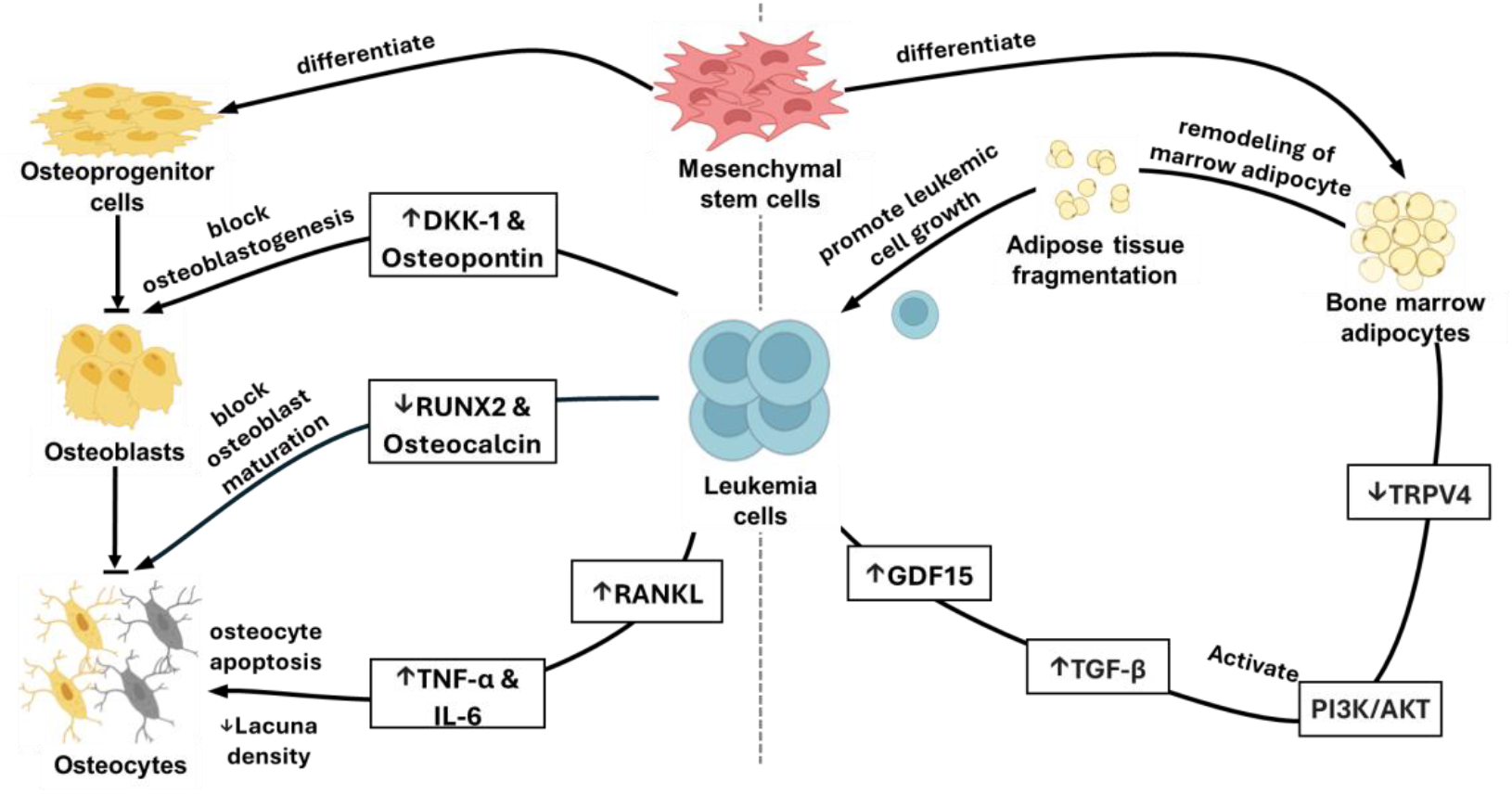
Proposed mechanism of CLL-induced disruption in bone marrow microenvironments. Leukemic cells (left) may inhibit osteoblastogenesis by increasing DKK-1 and osteopontin, reducing RUNX2 and osteocalcin, and promoting osteocyte apoptosis via elevated TNF-*α* and IL-6 through RANKL signaling, resulting in reduced osteocyte lacunar density. Leukemic cells (right) may induce adipocyte fragmentation by secreting GDF15 and TGF-*β*, activating PI3K/AKT, and suppressing TRPV4, creating a metabolic feedback loop supporting leukemic growth.

Lacuna aspect ratio and adipose density did not differ significantly between groups. However, adipose density demonstrated a noteworthy inverse correlation with lacunar density (r = –0.60), further emphasizing the interplay between bone and adipose compartments. Peak tissue mineral density similarly displayed no significant groupwise differences but correlated moderately with median lacuna volume (r = 0.50), indicating that local changes in mineralization are occurring in parallel with lacunar enlargement rather than a uniform global shift in bone mineral content.

Further analysis of lacunar morphology revealed a nuanced picture. Although a comparison of patient-level medians showed that lacunar volume was not statistically different between the groups (p=0.21), a more sensitive Kolmogorov-Smirnov test on the entire population of individual lacunae revealed that the overall distributions of both lacunar volume and aspect ratio were, in fact, highly significantly different (p ¡ 0.001 for both). This indicates that while the central tendency of lacunar size is only slightly shifted (an approximate 12% increase in the Hi-CLL median), high CLL burden induces more complex and significant alterations in the heterogeneity and shape profile of the lacunar network.

Collectively, these findings underscore the complex and multifaceted nature of CLL-associated changes in the marrow microenvironment, emphasizing the potential biological significance of lacunar enlargement and its relationship to adipose tissue disruption and osteocyte density reduction.

## Discussion

Our finding of a 40.3% reduction in osteocyte lacunar density with high CLL involvement is particularly critical, given that osteocytes are the primary regulators of skeletal homeostasis. As the most abundant cells in bone, they are responsible for orchestrating remodeling and sensing mechanical load [31]. A decreased density of osteocyte lacunae implies fewer functional osteocytes, potentially compromising bone strength, remodeling efficiency, and fracture resistance [32]. Clinically, this could partially explain the elevated fracture risk reported in CLL patients, despite seemingly normal or borderline DXA bone density scores [6].

Several converging lines of evidence indicate that the decline in osteocyte lacuna density observed in CLL originates upstream, at the level of osteoblast maturation. Under physiologic conditions, committed osteoblasts lay down mineralized matrix and, upon full maturation, become embedded as osteocytes. Giannoni et al. [33] demonstrated that co-culture of bone-marrow stromal cells with CLL cells halts this process: key markers of mature osteoblasts (RUNX2, osteocalcin) fall sharply, and inhibitors of osteoblastogenesis (DKK-1, osteopontin) rise. Mechanistically, CLL cells secrete high levels of TNF-*α*, IL-6, and IL-11; neutralizing these cytokines or silencing their genes in CLL cells restores normal mineralization, indicating a potential causal link. Because new osteocytes can form only after successful osteoblast mineralization and embedding, a cytokine-driven arrest of osteoblast maturation, as demonstrated in other studies, could explain the reduction in osteocytes incorporated into bone, thereby lowering overall lacunar density observed in our data.

Osteocyte apoptosis represents another critical pathway contributing to reduced lacuna density. Chronic inflammatory environments typical of CLL are rich in pro-inflammatory cytokines such as TNF-*α* and IL-6 [34, 35], both of which have been documented to induce osteocyte apoptosis in vitro [36, 37]. Increased osteocyte apoptosis subsequently amplifies bone resorption activity through mechanisms such as RANKL release, further disrupting the integrity and functionality of bone.

Analogous findings from other hematologic malignancies provide additional context for interpreting our results. MM [14] extensively disrupts the bone marrow microenvironment through mechanisms involving enhanced suppression of osteoblast differentiation and osteocyte apoptosis. Although typically less aggressive, our findings indicate that CLL similarly induces subtler, yet measurable, disruptions of the osteocyte network. In contrast to MM where treatment is recommended immediately at diagnosis, observation instead of active treatment is recommended in a large population of patients with CLL, which could entail years where the disruption of the bone marrow microenvironment can continue uninhibited by CLL therapy. Overall, the parallels in remodeling of the bone marrow microenvironment across different hematologic malignancies underline common mechanistic themes, potentially suggesting that targeted interventions effective in MM could be explored for therapeutic potential in managing skeletal health in CLL.

Our observation of increased adipose tissue fragmentation, as indicated by elevated adipose tissue SA:V, provides novel insights into the bone marrow microenvironment alterations associated with CLL infiltration. Despite similar BMI between lo-CLL and Hi-CLL groups (28.7 vs 29.3; Table 1), these microstructural differences indicate that the adipose changes are not explained by systemic adiposity alone. Taken together, the findings are consistent with local leukemia–adipocyte interactions and niche inflammation contributing to adipose fragmentation, although causal mechanisms remain to be established and will require targeted mechanistic studies.

While the specific pathways in CLL require further investigation, studies in other leukemias offer a potential mechanism. Patients with CLL have elevated peripheral blood levels of Growth Differentiation Factor 15 (GDF15) [38], a factor shown in AML to promote adipocyte remodeling through complex downstream signaling [39]. In AML models, leukemia-derived GDF15 and stromal-derived TGF-*β* have been shown to co-activate the PI3K/AKT signaling pathway in adipocytes [40]. If a similar pathway is active in CLL, it could explain the observed structural changes, as enhanced PI3K/AKT signaling is known to suppress the transcription factor FOXC1, a key regulator of adipocyte integrity. The downstream effect of reduced FOXC1 activity in these models is the downregulation of TRPV4, a channel protein essential for maintaining normal adipocyte morphology and volume. This can lead to a phenotype of smaller, remodeled adipocytes with altered metabolic properties, including enhanced lipolysis [40]. Such a process could create a metabolically favorable niche that provides energy substrates to fuel leukemia cell growth.

This adipocyte remodeling mirrors observations in other hematologic malignancies where altered adipocyte metabolism supports malignant cells [41]. Our results suggest that CLL, despite its typically indolent course, may employ analogous metabolic reprogramming mechanisms within the marrow to support leukemic cell survival. The structural fragmentation we identified is a key piece of evidence consistent with this hypothesis. Future studies targeting pathways such as GDF15 or PI3K/AKT signaling could therefore offer novel therapeutic approaches for mitigating marrow adipose tissue disruption in patients with CLL.

Our findings reveal a significant negative correlation between osteocyte lacuna density and adipose fragmentation, highlighting a previously unrecognized interplay between the bone and adipose compartments in CLL. This relationship suggests that leukemic infiltration is associated with simultaneous and coordinated changes in both skeletal integrity and adipose remodeling.

This observation is contextualized by the well-established crosstalk between bone and adipose tissues, which is critical for marrow homeostasis [42, 43]. This balance is reportedly disrupted in other malignancies where leukemic cells can skew mesenchymal stem cell (MSC) differentiation away from functional osteoblasts and towards dysfunctional adipogenesis [44]. The concurrent loss of osteocytes and adipose fragmentation we observed could plausibly reflect a similar skewed differentiation process. Furthermore, since osteocytes and adipocytes mutually regulate each other through secreted factors [45, 46], the depletion of osteocytes in CLL may further impair adipocyte function, creating a detrimental feedback loop.

This pattern of interconnected remodeling is consistent with findings in other hematologic cancers. For instance, both AML and MM are known to reprogram marrow adipocytes to create a pro-tumorigenic niche that promotes disease progression and contributes to bone destruction [47, 48]. Our data suggest that CLL may induce a similar, complex interplay that disrupts the marrow microenvironment, thereby supporting disease progression.

This study has some limitations. First, there was no non-CLL control group; all bone marrow samples analyzed were obtained from patients already diagnosed with CLL because bone marrow core biopsies are invasive procedures that are not ethically or routinely performed in healthy individuals. This limits our ability to compare findings to a baseline of healthy bone marrow microarchitecture and underestimates the full extent of CLL-associated changes. In addition, while SRµCT provided detailed structural insights, complementary histological and biochemical analyses were not performed, leaving proposed mechanisms such as osteocyte apoptosis or cytokine-mediated remodeling unevaluated at the cellular level.

Future research should aim to address these limitations. Incorporating histological assessments of osteocyte viability and apoptosis markers, alongside cytokine profiling, would enhance mechanistic understanding. Exploring targeted therapeutic interventions to preserve osteocyte function and prevent adipose tissue fragmentation should also be prioritized in subsequent research. Such investigations could ultimately inform clinical practice by providing this mechanistic understanding while also bridging the bone quality indicators investigated using SRµCT to more accessible microscopy imaging.

Our findings have important clinical implications, primarily by identifying potential new biomarkers for assessing skeletal health in CLL. Current tools like DXA scans are often insufficient for predicting the high risk of fragility fractures in these patients. The microstructural changes we identified, osteocyte loss and adipose fragmentation, could serve as more sensitive indicators of marrow deterioration. While SRµCT is a research tool, these features could potentially be adapted for evaluation in standard clinical pathology workflows, for instance, through quantitative histology of bone marrow biopsies.

Furthermore, the observed interplay between bone and adipose tissue highlights novel therapeutic avenues, such as targeting metabolic pathways in adipocytes or using cytokine neutralization to preserve osteocyte viability. Critically, our results provide the first quantitative evidence that uninhibited leukemic growth actively degrades bone quality, offering a compelling microstructural explanation for skeletal fragility in this disease. This finding directly challenges the standard “watch and wait” approach for asymptomatic CLL, raising the important question of whether earlier therapeutic intervention is necessary to prevent the irreversible skeletal complications that cause significant morbidity in patients. Ultimately, our work suggests that the preservation of skeletal health should be a more central consideration in the future management of CLL.

## Acknowledgments

The authors want to greatly thank the patients who contributed bone marrow biopsy samples to this research. All bone marrow cores were obtained after patients provided informed consent and approved by the institutional review board (IRB# 00045880) from the University of Utah Huntsman Cancer Institute. We acknowledge support from the University of Utah Huntsman Cancer Institute for bone marrow biopsy collection. Also, We also thank the following team members from Total Cancer Care at Huntsman Cancer Institute for obtaining written consent from patients and for collecting bone marrow cores:Emily Simons, Paige Ward, Cailey Hohman, Sharmilee Nuli, Macy Barrios, Sydney Loveless, Natalie Rojas, Erica Wolff, Allanah Beazley, B. Stephanie Simmons, Andrea Campos, Brady Groneman, Nic Siniscalchi, Morgan Marshall. This work used resources from the Advanced Light Source at beamline 8.3.2., a U.S. DOE Office of Science User Facility under contract no. DE-AC02-05CH11231. We thank Dilworth Y. Parkinson for assistance in data acquisition.

In Kyu Lee was supported by the Department of Mechanical and Aerospace Engineering Fellowship at the University of California, San Diego. Deborah M. Stephens was supported by the NIH National Cancer Institute (grant R50CA275929).

