## Supplementary figures and images for "Disruption of Marrow Microenvironments in Chronic Lymphocytic Leukemia by High-Resolution Synchrotron Micro-Computed Tomography"

### Supplemental figure 1

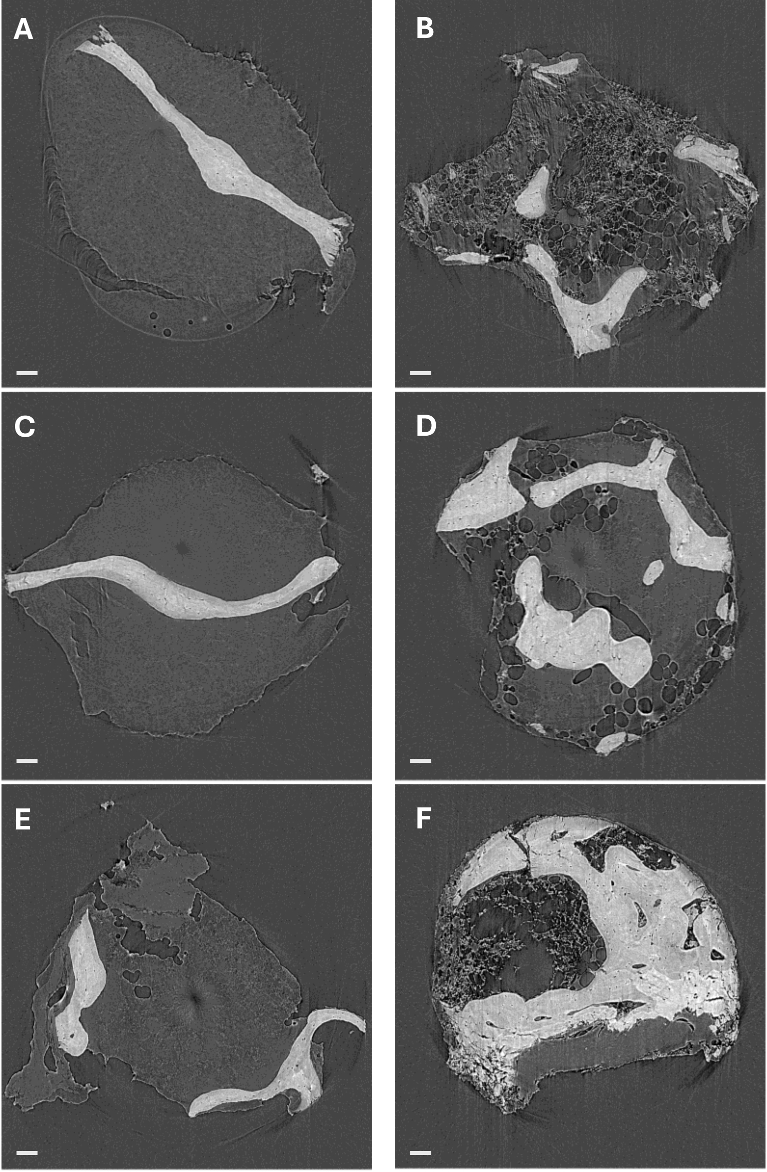
